# Transient upregulation of translational efficiency in prodromal Tg2576 mice precipitates AD symptoms

**DOI:** 10.1101/581652

**Authors:** Antonella Borreca, Francesco Valeri, Mariassunta De Luca, Lysianne Ernst, Arianna Russo, Alberto Cordella, Veronica Corsetti, Annalisa Nobili, Giusy Amadoro, Nicola Biagio Mercuri, Marcello D’Amelio, Martine Ammassari-Teule

**Affiliations:** CNR-IBCN, Via Del Fosso di Fiorano, 64. Rome; IRCCS Fondazione Santa Lucia, Centro Europeo di Ricerca sul Cervello (CERC), via del Fosso di Fiorano, 64, 00143, Rome, Italy; University of Namur, Faculty of Sciences, Belgium; INGM National Institute of Molecular Genetic, Via Francesco Sforza, 35–20122 Milano, Italia; Unit of Molecular Neurosciences, Department of Medicine, University Campus-Biomedico, via Alvaro del Portillo, 21 00128 Rome, Italy; EBRI FOUNDATION Viale Regina Elena 295, Roma; IFT-CNR Via Fosso del Cavaliere 100 - 00133 Roma; University of Rome “Tor Vergata”, 00133 Rome Italy

**Keywords:** Alzheimer Disease, Tg2576 mice, hAPP mRNA, eIF2α, translational control, salubrinal, pre-symptomatic markers, cognition, synaptic plasticity

## Abstract

Tg2576 mice overexpressing APPK670/671L exhibit elevated hAPP levels from birth but remain at a prodromal stage until 3 months of age. Whether variations in hAPP mRNA specific and overall translation occur during development and precipitates the transition from an asymptomatic to a symptomatic stage is unknown. By performing polysome profiling and distribution of hAPP, and measuring the levels of eukaryotic initial translation factors in hippocampal extracts from pre- and early symptomatic Tg2576 mice, we found that the presence of mRNA and protein polysomal signals was associated with decreased levels of the phosphorylated form of the initial translation factor eIF2α (p-eIF2α), revealing a transient upregulation of overall translation. Differently, the reduction of hAPP mRNA polysomal signals was associated with increased p-eIF2α levels – repressing translation-when mice were symptomatic, suggesting a compensatory mechanism aimed at downregulating hAPP mRNA. Confirming that prodromal upregulation of translational efficiency contributes to AD pathogenesis, pharmacological restoration of proper translational control in early symptomatic mice blocked the manifestation of neural and cognitive AD-like alterations.

## Introduction

In mouse models of AD, overexpression of mutant forms of the human amyloid precursor protein (hAPP) promotes formation of toxic Aβ and impairs cognition at age points which vary according to the locus of the mutation in the APP gene and the number of hAPP transgene copies (Howlett DR et al., 2009). Among mice carrying the Swedish APP KM670/671N mutation, there is evidence that the Tg2576 model develops normally until 3 months of age. Then, cognition progressively declines concurrently with the augmentation of Aβ load until the manifestation of a full AD symptomatology. Although the insertion of the hAPP transgene in the mouse genome warrants that hAPP is overexpressed from birth in this model, little is known as whether variations in the levels of the protein and of its messenger occur during development and precipitate the transition from the prodromal to the symptomatic stage.

We previously started to investigate this point by measuring hAPP mRNA and protein levels in hippocampal extracts from Tg2576 mice at three stages of development, and found that these levels considerably vary according to the mouse age (Borreca A et al., 2016). They are maximal when mice are asymptomatic (1 month of age), remain elevated at the onset of AD symptoms (3 months of age), and then decrease in association with the full symptomatology (6 months of age). Remarkably, pre-symptomatic maximal upregulation of hAPP was found to occur in association with the stronger unbalance of the RNA-binding proteins hnRNPC and FMRP, i.e., two post-translational opposite regulators of hAPP expression (Lee EK et al., 2010) which, being respectively up- and down-regulated, boost hAPP mRNA translation. Of note, we detected a similar enhancement of hAPP mRNA and protein with the same opposite dysregulation of FMRP and hnRNP C in hippocampal extracts from sporadic AD patients. These findings reveal that hAPP translation is persistently upregulated independently from mutations in the hAPP gene, and suggest that repression of translation should be beneficial for preventing/rescuing AD.

Consistent with this view, compounds known to reduce overall translation have been reported to decrease full length APP and to reduce Aβ deposition in a variety of *in vitro* preparations (Asuni AA et al., 2014), and in the brain of fully symptomatic hAPP mutant mice which otherwise showed a rescue neural plasticity and cognitive alterations (Asuni AA et al., 2014; Teich AF et al., 2017).

Although these findings show a regulatory role of overall translation on hAPP levels, how specific *vs* general translation are dysregulated in Tg2576 mice during development and contribute to age-specific variations in translational efficiency is unknown. To investigate this point, we performed polysome gradient analyses to identify the distribution profile of hAPP mRNA and protein in hippocampal extracts from pre- and early symptomatic mice, and from symptomatic mice and sporadic AD patients. In the same samples, we controlled overall translation by measuring the levels of eukaryotic initial translation factors. Revealing that specific and overall translation are concurrently upregulated at prodromal and early symptomatic stages, polysomal hAPP mRNA and protein distribution were found to be associated with decreased levels of the phosphorylated form of the initial translation factor eIF2α (p-eIF2α) in 1- and 3-month old mutants. Conversely, still elevated hAPP levels no longer associated with hAPP mRNA polysomal signals, but with enhanced peIF2α levels downregulating overall translation, were detected in symptomatic mice and symptomatic AD patients, suggesting a compensatory mechanisms aimed at downregulating hAPP levels. Consistent with a role for the early upregulation of translational efficiency in triggering AD pathogenesis, pharmacological restoration of proper translational control in early symptomatic mice prevented the manifestation of neural and cognitive alterations.

## METHODS

### Animals

Male mice overexpressing the APP695 fragment with the Swedish mutation subsequently backcrossed to C57BL/6 × SJL F1 females. The offspring was genotyped to confirm the presence of human mutant APP DNA sequence by PCR. Genomic DNA isolated from mouse tails was genotyped by polymerase chain reaction (PCR) analysis. Specific primers sense (5’-CTG ACC ACT CGA CCA GGT TCT GGG T-3’) and antisense (5’-GTG GAT AAC CCC TCC CCC AGC CTA GAC CA-3’) of APPSWE (10 μM each) were added into genomic DNA template mixtures that were subjected to 25 cycles of amplification. Amplification was conducted in a T100 thermal cycler (BioRad Laboratories Inc., Hercules, California, USA) under the following conditions: denaturation for 30 s at 94°C, annealing for 30 s at 62°C, and extension for 45 s at 72°C. The amplified PCR products were then loaded onto a 1.0% agarose gel, after which the bands were detected using the Kodak Electrophoresis Documentation and Analysis System 120 (Eastman Kodak, Rochester, NY, USA). Mice were reared maintained on a 12 h light/dark cycle with ad libitum access to food and water. All experiments were performed in accordance with the guidelines provided by the European Communities Council Directive of 24 November 1986 (86/609/EEC).

### Polyribosome profile

Polyribosome profiling allows to study the overall degree of translation of individual proteins and their mRNAs. Hippocampi from Tg2576 mice of 1, 3, and 6 months of age (pool of 4 animals for each age point) and human brain post mortem were homogenized in 0,5 ml of lysis buffer (100 mM NaCl, 10 mM MgCl_2_, 10 mM Tris-HCl pH 7.5, 1% Triton-X100). The lysates were incubated 5 min in ice, centrifuged for 5 min at 12,000 g at 4°C, and the total amount of supernatants (cytoplasmic extract) centrifuged through 15%–50% (w/v) sucrose gradients for 180 min at 37,000 rpm in a Beckman SW41 rotor. The polysomes, the 80S monosome, the two subunits 60S and 40S as well as the very light mRNPs were detected by UV absorbance at 256 nm (BioRad) and each gradient was collected in 10 fractions. From each fraction, the proteins were precipitated with mix containing 50% Ethanol, 25% Methanol and 25% Acetone. The proteins extracted were resuspended in 30 μl of laemly buffer 2x and then analyzed by Western blotting.

### Western blot determination of APP on polysome fractions

From each polysomal fraction, we used 200 μl for protein extraction. 200 μl of each fraction was treated with 6 volumes (1,2 ml) of mix containing 50% EtOH, 25% MetOH and 25% Acetone and left overnight a 4°C. The day after each fraction was centrifuged at 14000g for 40 minutes and washed with 1 ml of EtOH 80%. The resulting pellet was resuspend in 30 μl of laemly buffer 2x. 10 μl of each fraction of polysomes gradient from WT and Tg2576 mice were used in this analysis. Total proteins were separated by 4-15% gradient sodium dodecyl sulfate–polyacrylamide gels (BIO-RAD Laboratories, Hercules, CA) and transferred to nitrocellulose membrane (BIO-RAD Laboratories, Hercules, CA). Western blots were blocked in 5% non-fat dry milk in TBST buffer (0.1% Tween 20 in Tris–borate saline) then incubated with APP (1:1000; Sigma). Blots were then incubated with appropriate conjugated to horseradish peroxidase (Chemicon) and developed by ECL Western Blotting Analysis System (GE Healthcare).

### Real Time-quantitative PCR for APP mRNA on polysome fractions

Cytoplasmic extracts prepared from hippocampi of Tg2576 mice were fractionated through sucrose gradients. Ten fractions were collected from each gradient while recording the absorbance profile. For RNA purification, the fractions were immediately treated with a solution containing 1% SDS (final concentration), 10 μg of glycogen, 50 pg of kanamycin positive control RNA and 100 μg/ml of proteinase K, and then incubated for 30 minutes at 37°C. Kanamycin synthetic RNAs were used to normalize for possible RNA loss during RNA phenol/chloroform extraction and precipitation from each fraction. RNAs were precipitated with 0.2 M NaOAc pH 4.8 and 0.7 volumes of isopropanol. The pellets, washed with EtOH 80%, were re-suspended in 20 µl of ddH_2_O. Total RNA extracted from each fractions was analyzed by quantitative RT-PCR for specific mRNAs (human APP). To correct for variations in the efficiency of the RT-PCR reaction, the same amount of a synthetic RNA (Kanamicin) was added to each sample, amplified, and used for normalization. 1 μl of each fraction were pulled to create polysomes fractions (Fraction 1-5) and non polysomes fraction (Fraction 6-10). 5 μl of each fraction (polysomes and non polysomes) was used for First-strand synthesis using p(dN)6 and 100 U of M-MLV RT (Invitrogen). The same amount of first strand DNA was used for qRT-PCR analysis. The qRT-PCR was conducted with SYBR green master mix (Applied Biosystem) and specific primer for APP mRNA and Kanamycin synthetic DNA.

### Western blot determination of eukaryotic initial translation factors

Hippocampi from WT and Tg2576 mice were collected at each age-point of interest and homogenized in RIPA buffer (10 mM Tris-HCl, pH 7.5, 150 mM NaCl, 2% Nonidet P-40, 5 mM EDTA, 0.1 mM phenylmethylsulfonyl fluoride, 1 mM β-glycerophosphate, 1 mM sodium orthovanadate, 10 mM sodium fluoride, 0.1 M SDS, 1% protease inhibitor cocktail-Sigma Aldrich) (Borreca A et al., 2018). A pool of 4 animals was used for the analysis. To confirm these data a four WT and Tg2576 mice were loaded on acrylamide gel separately. 50 μg of proteins were loaded on 12% acrylammide gel and transferred on nitrocellulose membrane, blocked with 5% not fat milk and incubated with p-eIF2α and eIF2α antibodies (Cell signaling) over night at 4°C (diluted in BSA 5%). After three washes with TBST, the membranes were incubated with secondary antibody HRP, specific for both primary antibodies and the signal was revealed by ECL Western Blotting Analysis System (GE Healthcare). Additional p-eIF4E, eIF4E and eIF4G antibodies (Cell signaling) were used to determine the level of these two other major eukaryotic initial translation factors in the hippocampus of 3-month old Tg2576 mice.

### Pharmacological treatment

Pharmacological inhibition of elF2α (eukaryotic translation initiation factor 2 subunit alpha) dephosphorylation was carried out by injecting salubrinal (Tocris Cat. No. 2347) emulsified in DMSO (Sigma) (1% diluted in saline). Two administration routes were used according to previously established injection protocols (Sokka AL et al., 2007). Mice were given either one intra-cerebro-ventricular (i.c.v.) injection (1 μl of a 75 μM solution) or one daily intraperitoneal (i.p.) injection repeated for 7 consecutive days (1 mg/kg). Control mice received i.c.v or i.p. injections of DMSO.

### Western blot determination of APP, Aβ, and BACE-1 levels in hippocampal extracts

Hippocampi of WT and Tg2576 mice, treated with salubrinal or DMSO, were lysed with RIPA buffer (10 mM Tris-HCl, pH 7.5, 150 mM NaCl, 2% Nonidet P-40, 5 mM EDTA, 0.1 mM phenylmethylsulfonyl fluoride, 1 mM β-glycerophosphate, 1 mM sodium orthovanadate, 10 mM sodium fluoride, 0.1 M SDS, 1% protease inhibitor cocktail-Sigma Aldrich)(Borreca A et al., 2016). After homogenization, samples were centrifuged at 12000 g for 10 minutes and the supernatant was collected, quantified with Bradford assay, and 50 μg of total protein were loaded on 12% acrylamide gel for determination of APP (Sigma), Amyloid-β (Aβ) peptide species (D54D2 Cell Signaling), β-secretase enzyme-1 (BACE-1, Millipore)

### Western blot for determination of Caspase-3 activity in hipppocampal synaptosomes

Hippocampi of WT and Tg2576 mice treated with salubrinal or DMSO were lysed with Homogenization buffer (320 mM sucrose, 4 mM Hepes pH 7.4, 1mM EGTA, 1 mM PMSF and 1x protease inhibitor cocktail). The homogenized tissue was centrifuged at 1000g for 10 minutes a 4°C. the supernatant was collected and centrifuged at 12000g for 15 minutes at 4°C. The obtained pellet was resuspended in homogenization buffer and centrifuge at 13000g for 15 minutes at 4°C. The pellet was resuspended in RIPA buffer (10 mM Tris-HCl, pH 7.5, 150 mM NaCl, 2% Nonidet P-40, 5 mM EDTA, 0.1 mM phenylmethylsulfonyl fluoride, 1 mM β-glycerophosphate, 1 mM sodium orthovanadate, 10 mM sodium fluoride, 0.1 M SDS, 1x protease inhibitor cocktail-Sigma Aldrich). The resuspended pellets was sonicated and centrifuged at 12000 rpm for 10 minutes at 4°C. The synaptosomes were loaded on acrylammide gel and the CASP3 (Cell signaling) levels (cleaved/total) (D’Amelio M et al., 2011) was evaluated.

### Acute slices preparation for electrophysiology

Mice were deeply anesthetized by inhalation of 2-Bromo-2-Chloro-1,1,1-trifluoroethane and after decapitation the brains were rapidly removed from the skull. Parasagittal slices were cut with a vibratome (VT1200S, Leica) and then immersed in chilled bubbled (95% O2, 5% CO2) aCSF containing (in mM): NaCl 124, KCl 3, NaH_2_PO_4_ 1.25, NaHCO_3_ 26, MgCl_2_ 1, CaCl_2_ 2, glucose 10 (∼ 290 mOsm, pH 7.4). The slices were incubated for 1 h in CSF at 32°C and then left at room temperature for 30 min before recordings.

### Field recordings of excitatory postsynaptic potentials

A single brain slice was transferred to a recording chamber of an upright microscope (Axioskop 2-FS; Zeiss, Germany) and completely submerged in aCSF (3–4 ml/min; 32 °C). Field excitatory postsynaptic potential (fEPSP) were induced by stimulation (100 µs duration; every 30 s) of Schaffer collateral pathway using a concentric bipolar stimulating electrode (FHC Inc.; Bowdoin, ME) and a CSF-filled borosilicate glass recording electrode that were positioned in the stratum radiatum of the CA1 hippocampal region at a distance of 200-300 μm. All experiments were performed at the intensity yielding a half-maximal response of input-output curves that was obtained by measuring the fEPSP initial slope at increasing 10 µA steps of afferent stimulation (D’Amelio M et al., 2011).

### Long Term Depression

LTD was induced 20 min after the test stimulation (at half-maximal intensity, every 30 s) when fEPSP slope stability was obtained. The slice was challenged with DHPG (50 μM) for 10 min and then washed out for 60 min. The DHPG-LTD was evaluated by the fEPSP mean slope 55-60 min from DHPG washout, normalized to the mean slope during baseline, and recorded during the 10 min preceding DHPG perfusion. The field responses were recorded with a MultiClamp 700B amplifier and digitized with Digidata 1322A. Data were sampled at 20 kHz. Traces were obtained by pClamp 9.2 and analyzed using Clampfit 9.2 (all from Molecular Devices; Sunnyvale, CA).

### Golgi staining

After one week of pharmacological treatment with salubrinal or DMSO, mice were deeply anaesthetized with a cocktail of Zoletil (800mg/kg) and Rompum (200mg/kg) and perfused transcardially with 0.9% saline solution (*N* = 7 mice per group). Brains were dissected and immediately immersed in a Golgi-Cox solution (1% potassium dichromate, 1% mercuric chloride, and 0.8% potassium chromate) at room temperature for 6 days according to a previously described protocol (Gibb R et al., 1998). On the seventh day, brains were transferred in a 30% sucrose solution for cryoprotection and then sectioned with a vibratome. Coronal sections (100 μm) were collected and stained according to the method described by Gibb and Kolb (1998). Sections were stained through consecutive steps in water (1 minute), ammonium hydroxide (30 minutes), water (1 minute), developer solution (Kodak fix 100%, 30 minutes), and water (1 minute). Sections were then dehydrated through successive steps in alcohol at rising concentrations (50%, 75%, 95%, and 100%) before being closed with slide cover slips. Spine density was analyzed on CA1 neurons. Neurons were identified with a light microscope (Leica DMLB) under low magnification (20×/NA 0.5). Five neurons within each hemisphere were taken from each animal. On each neuron, five 30– 100 μm dendritic segments of secondary and tertiary branch order of CA1 dendrites were randomly selected and counted using Neurolucida software. Only protrusions with a clear connection of the head of the spine to the shaft of the dendrite were counted as spines. Statistical comparisons were made on single neuron values obtained by averaging the number of spines counted on segments of the same neuron. The analysis was conducted by an experimenter blind to the experimental condition.

### Novel object recognition

The novel object recognition (NOR) test was carried out in 3-month old Tg2576 and WT mice. Testing started on the day after mice received the last i.p. injections of salubrinal or DMSO. Mice in their home cage were transferred to experimental room and left to acclimate for 1 h to the new environment. NOR testing consisted in three sessions (Fig. 5A). On the first session (open field exploration), each mouse was placed in an empty squared open field (40 cm in side) surrounded by 60 cm-high walls and left free to explore it for 10 minutes. The mouse was returned to its home cage for a 10-min pause during which two identical glass cylinders of 3 cm in diameter and 10 cm in height (object on the left: sx; object on the right: dx) were put in opposite corners of the open field. On the second session (training), the mouse was placed in the center of the open field and allowed to explore objects sx and dx for 10 minutes. The mouse was returned again to its home cage for a 1 h-pause during which one (sx or dx) familiar object (FO) was substituted with a novel object (NO), a multicolored kubrik cube of 5 cm in side (NO). On the third session (testing), the mouse was placed again the center of the open field and allowed to explore the FO and the NO for 10 min. Object exploration was defined as mice sniffing or touching the object with its nose and/or forepaws. The objects were cleaned with 10% ethanol between each session. The preference index was calculated according the formula previously described (Antunes M et al., 2012) which estimates the percentage of time spent exploring each object (FO or NO) over the total time spent exploring both objects. A preference index above 50% indicates a preference for the NO, below 50% a preference for the FO, and 50% no preference.

**Figure 5.**
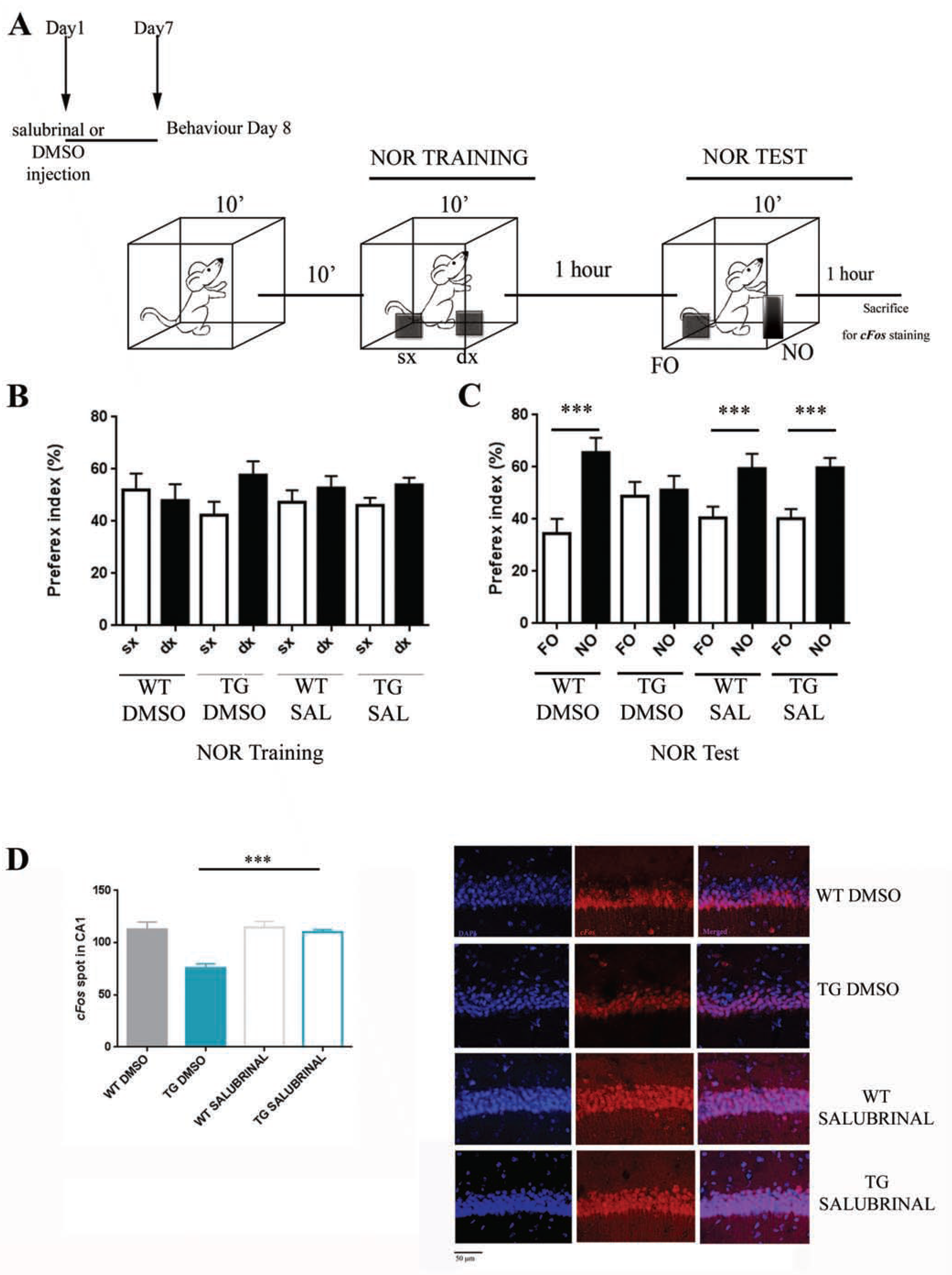
Salubrinal rescues novel object recognition deficit in Tg2576 mice (A) Cartoon depicting the behavioral protocol. Tg2576 mice were injected daily with salubrinal (i.p., 1 mg/kg) or with the same volume of DMSO during 7 consecutive days and their performance in the novel object recognition (NOR) task was compared with the performance of wild-type (WT) mice injected with DMSO. Mice were first exposed to two identical objects (training) and, 1 h after, one of familiar object was substituted with a novel object (testing). The time spent exploring each object was recorded during each phase and the preference index for each object was calculated (time exploring one object divided by the time exploring the two objects*100). (**B)** Histograms showing the preference index for object sx (left, white bars) and dx (right, black bars) during training. No effect of genotype or treatment was found (**C**) Histograms showing the preference index for object FO (familiar object) and object NO (novel object). Tg2576 mice injected with DMSO showed the same preference index for objects FO and NO. Differently, Tg2576 mice injected with salubrinal, and wild-type mice injected with DMSO explored significantly more object NO than object FO. (**D**) Histograms (left) and representative c-fos immunohistochemistry staining (right) showing the number of c-fos positive spots in the CA1 region of the hippocampus (10 areas of 25×25 μm in size per mouse) of DMSO-injected wild-type (grey solid bars) and Tg2576 (blue solid bars), and in salubrinal injected wild-type (grey empty bars) and Tg2576 (blue empty bars) mice. Bars represent mean ± SEM. \**p* < 0,05.

**Figure 6:**
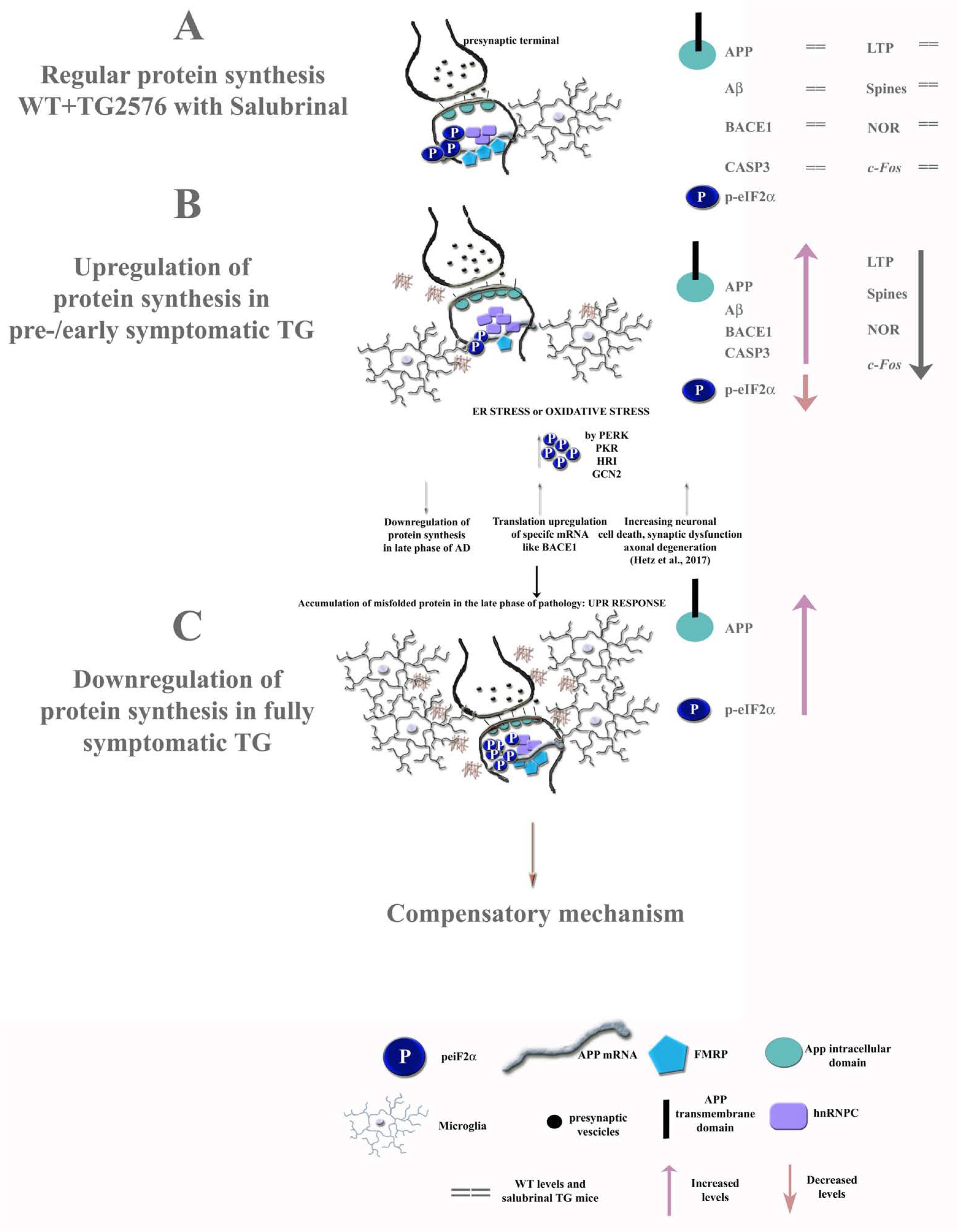
A cartoon depicting a possible molecular mechanism. In the early symptomatic phase of the pathology (**B**) the levels of p-eIF2α is reduced with increasing of protein synthesis machinery and with higher levels of APP protein. As a consequence, we observe an increase of Aβ production, activation of inflammatory response, ER stress o Oxidative stress induction with compensatory mechanism of downregulation of protein synthesis (**C**). On the other hand we observe a translation upregulation of specific mRNA (BACE1), an increase of neuronal cell death, synaptic dysfunction and axonal degeneration. The early AD symptoms are entirely rescued by pharmacological (salubrinal) downregulation of overall translation (**A**).

### NOR-induced *c-fos* activation

1 h following NOR testing, mice were deeply anesthetized with mix solution of ketamina/xylazin (100 mg/kg) and immediately perfused with PAF 4% according to a procedure previously described (Borreca A et al., 2018). The brains were immediately removed, post-fixed in PAF 4% overnight, transferred in sucrose (30% diluted in PBS1X), and then sectioned coronally with a cryostat (40 μm). Slices were immediately washed with PBS 1X and incubated with primary antibody (*c-fos* 1:400) diluted in a solution of PBS 1X and 0.3% Triton overnight at 4°C. After incubation, the slices were immediately washed with PBS 1X and incubated with specific secondary antibody (Alexa Fuor 555 antirabbit) for two hours at room temperature avoiding light. DAPI staining (1:1000, Enzo Life Science) was performed at the last wash with PBS 1X. Sections were mounted with fluoromount (Sigma) and coverslipped. The staining was visualized at confocal microscope (Zeiss LSM700; magnification 20X) and images were analyzed with IMARIS software. The number of *c-fos* immunoreactive spots in the CA1 hippocampus were counted in 10 areas of 25×25 μm were analyzed avoiding the DAPI. The analysis was conducted in the CA1 hippocampus. Number of spots for WT and Tg2576 mice were counted and statistical analysis were conducted.

### Statistical analyses

Group differences in hippocampal APP mRNA polysome gradient distribution and in p-eiF2α/eiF2α ratio were evaluated by means of two-way ANOVAs with genotype and age point as main factors. Student’s t-tests (two-tailed) for unpaired samples were used to evaluate group differences in p-eIF4E/eIF4E and eIF4G levels, and the effect of i.c.v. and i.p. injections of salubrinal on APP and Aβ hippocampal levels in 3-month old Tg2576 mice. In mice of the same age receiving DMSO or salubrinal injections, group differences in hippocampal levels of p-eIF2α, APP, BACE-1, cCASP-3/Casp-3 ratio, dendritic spines, LTD, and NOR-induced *c-fos* activation were statistically evaluated by means of two way ANOVAs with genotype and treatment as main factors. Student’s t-tests for paired samples were used to estimate object novelty preference in each experimental condition. Post-hoc pair comparisons were carried out where necessary by means of the Bonferroni and Fischer test. Statistical levels of significance were set at *P* < 0.05.

## RESULTS

### APP and APPmRNA polysomal signals were detected in pre-symptomatic and early-symptomatic Tg2576 mice

Polyribosome profiling sheds light on the mechanisms which regulate translation and protein synthesis. The integrity and distribution of polysomes were based on the UV profile (Fig. 1). Regarding protein expression, an APP signal was detected in both polysome (lanes 1-5) and non-polysome (lanes 6-10) fractions in 1-month old Tg2576 mice while it was present only in the non-polysome fraction in WT mice of the same age. A weaker signal was still detected in 3-month old Tg2576 mice, but was no longer present when mice were 6-month old. Thus, the presence of a polysomal APP signal is exclusively detected in Tg2576 mice and reaches its maximal expression at the age-point mice are cognitively asymptomatic (Fig. 1A). The L7 distribution was also evaluated (data not shown). Of note, also non-polysomal signals (lane 8-10) were decreased in both genotypes at 6 months of age.

**Figure 1.**
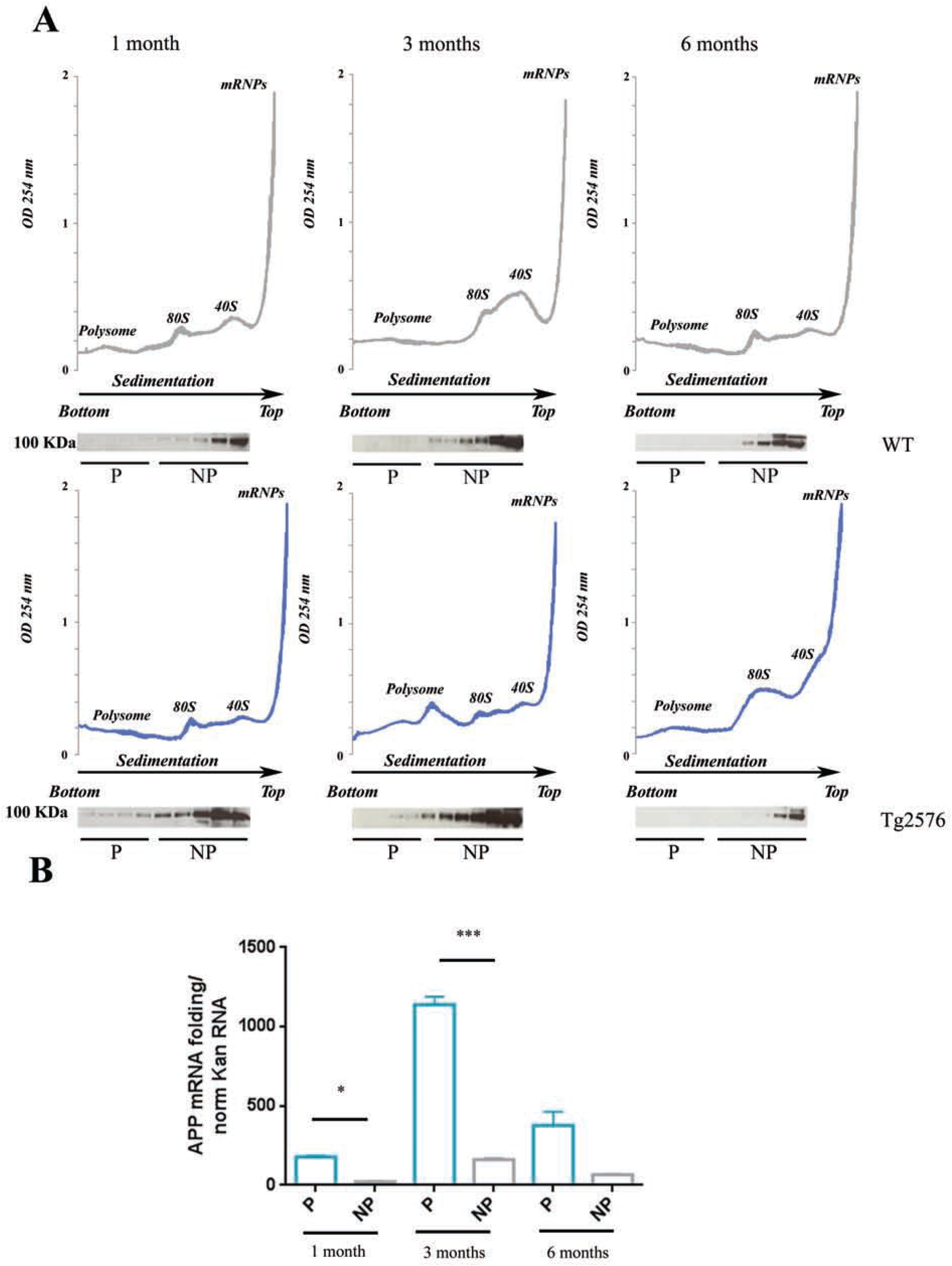
Polysomal gradient fractionation of APP protein and hAPP mRNA from hippocampal extracts isolated from wild-type and Tg2576 mice of 1, 3, and 6 months of age. Polysomal profile obtained from cytoplasmic brain extracts from wild-type mice (upper panels, grey lines) and Tg2576 mice (lower panels, blue lines). Brain extracts were centrifuged through a 15–50% sucrose gradient; absorbance at 254nm was monitored continuously and plotted against the fraction numbers. The bottom fraction (heavy) includes fractions 1–5 and corresponds to polysomes (P). The top fraction (light) includes fractions 6–10 and corresponds to non polysomes (NP) with 80S, 60S 40S and mRNPs particles. (B) Histograms showing the distribution of hAPP mRNA on P and NP fractions after further processing of total RNA from each fraction by RT– qPCR (normalization with the synthetic RNA Kanamicin). Bars represent mean ±SEM. The experiment was triplicated (n=4 mice for each condition in each experiment). \**p* < 0.05 for WT vs Tg2576.

A reliable way to assess mRNA translational efficiency is to analyze its partitioning between actively translating polysomes and mRNPs that are not translated. Total RNA was extracted from the gradient fractions and then analyzed by quantitative RT-PCR for hAPP mRNAs. Data are shown in Fig.1B and are expressed as levels of mRNA in polysomal vs non polysomal fraction (pool of four mice from each genotype reproduced three times). A two-way ANOVA performed on polysomal vs non polysomal APP mRNA levels detected in both genotypes at 3 age points revealed a significant genotype x age interaction (F_(2,12)_ = 326,42; *p* = 0,00001). Post hoc pair comparisons showed that APP mRNA was significantly more localized in the polysomal fraction when Tg2576 mice were 1-month (*p* = 0.0045) and 3-month (*p* = 0.0000001) old, but equally distributed in polysomal and non polysomal fractions when mice were 6-month old (*p* = 0.75), the latter observation being in line with data obtained in 9-month old mice bearing the triple APP/Tau/PS1 mutation (Caccamo A et al., 2015) and with our results in human AD brain post mortem (Suppl. Fig1). Thus, the major presence of APP mRNA in polysomal fractions is strictly associated with the upregulation of protein expression before or shortly after mice exhibit cognitive deterioration.

### Tg2576 mice show early upregulation and late downregulation of p-eIF2α

Synaptic dysfunction is the main cause of cognitive decline in AD. The eukaryotic initiation factor-2α (eIF2α) is essential for most forms of translation initiation and therefore for protein synthesis. In particular, its phosphorylation stabilizes the eIF-2/GDP/eIF-2B complex, prevents GDP/GTP exchange reaction, and produces global inhibition of translation. Extensive evidence suggests that aberrant phosphorylation of eukaryotic initiation factor-2α (p-eIF2α) induces synaptic failure and neurodegeneration through persistent inhibition of mRNA translation. In line with this hypothesis, Kim et al (2007) showed that p-eIF2α is upregulated in hippocampi homogenates from fully symptomatic 12-month-old Tg2576 mice. Interestingly, while we confirm that p-eIF2α is upregulated in hippocampal extracts from 6- and 9-month old symptomatic mice, here we show that this factor is downregulated at earlier stages of development (Fig. 2 A-B and Suppl Fig 2). A two-way ANOVA performed on p-eIF2α levels measured in both genotypes at four age points revealed a significant genotype x age point interaction (F_(3,16)_ = 77,40, *p* = 0.001). Subsequent post hoc pair comparisons then showed significant differences between genotypes at all age points developing, however, in opposite directions according to the age point. Compared to those detected in WT mice, p-eIF2α levels were decreased in Tg2576 mice at 1 month (*p* = 0.03) and 3 months (*p* = 0.0000001) of age but, consistent with previous data (Kim HS et al., 2007), these levels were increased when Tg2576 were 6-month (*p* = 0.000004) and 9-month (*p* = 0.02) old. These findings were confirmed by immunofluorescence detection of eIF2α phosphorylation expression in hippocampi sections taken from mice at the same age points (Fig. 2C). Of note, the non-phosphorylated form of eIF2α did not vary between genotypes in the asymptomatic (1 month: *p* = 0.79) and early symptomatic (3 month: *p* =0.18) phases, but was decreased in Tg2576 mice at 6 months (*p* = 0.04) and 9 months (*p* = 0.0005) of age, in association with the increase in the phosphorylated form. Altogether, these results point to the existence of non-linear variations of p-eIF2α levels in Tg2576 mice consistent with an early upregulation followed by a downregulation of the protein synthesis machinery which controls hippocampal APP expression.

**Figure 2.**
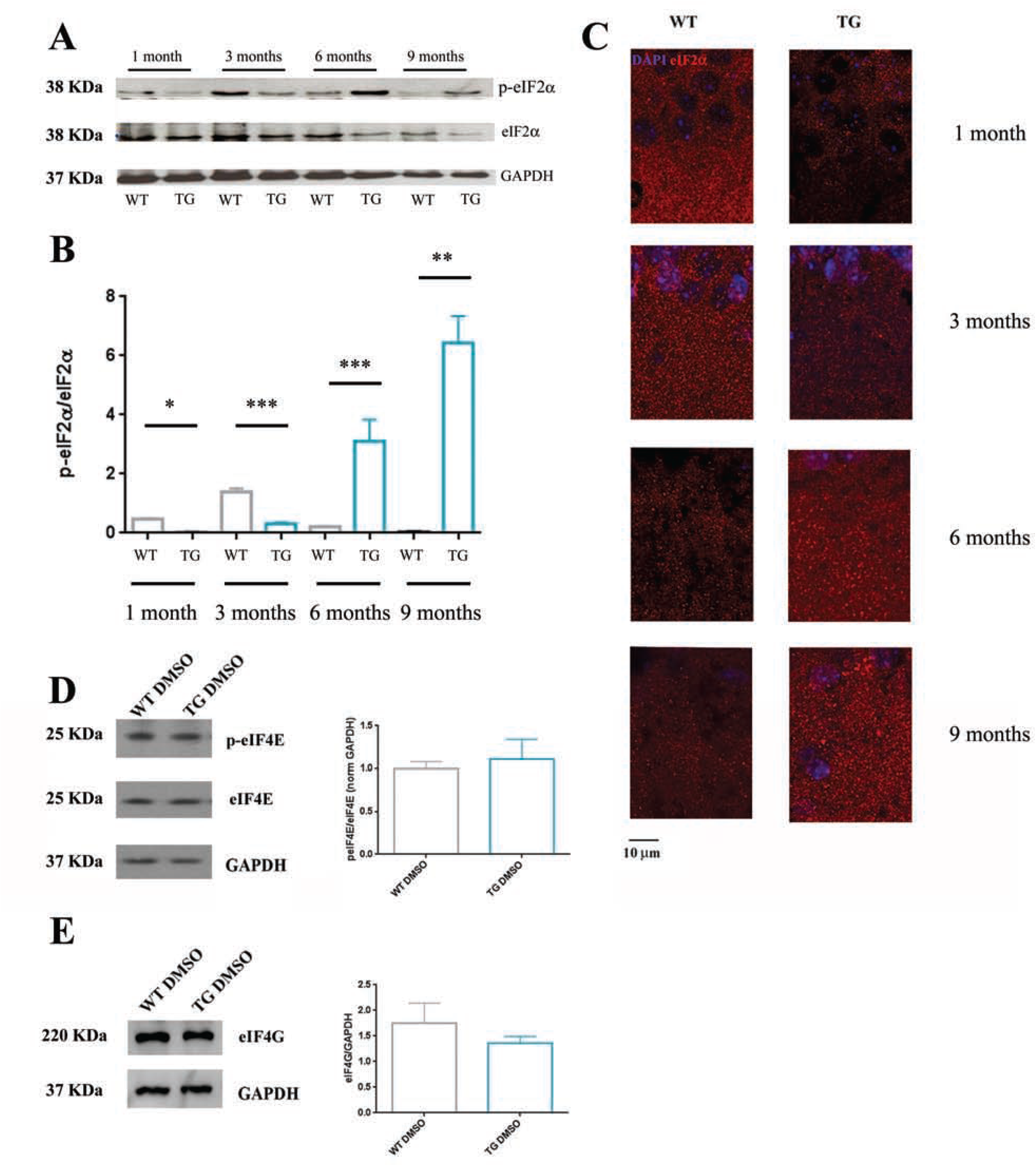
Hippocampal eIF2α levels in wild-type and Tg2576 mice across development. (**A**) Western blot analysis of phosphorylated and non-phosphorylated eIF2α levels normalized for GADPH in the hippocampus of wild-type mice and Tg2576 mice of 1, 3, 6, and 9 months of age. **(B)** Histograms showing variations of the peiF2α/eiF2α ratio in the hippocampus of the same mice groups. (**C**) Immunofluorescence visualization of hippocampal eiF2α levels in the same mice groups. The two analyses show that hippocampal eiF2α levels were decreased in pre-symptomatic (1 month) and early symptomatic (3 months) Tg2576 mice, but were increased in symptomatic (6 and 9 months) Tg2576 mice, compared to wild-type mice. *p* < 0,05 *; *p* < 0,01 **; *p* < 0,005 ***. Histograms showing **(D)** peIF4E/eIF4E and **(E)** eIF4G levels measured in the hippocampus of 3-month old wild-type and Tg2676 mice. No difference in the expression level of these two eukaryotic translation initiation factors was detected between genotypes. Bars represent mean + SEM. wild-type mice: grey bars; Tg2576 mice: blue bars.

### p-eIF4E/eIF4E and eIF4G are unaltered in early symptomatic Tg2576 mice

Eukaryotic translation initiation factor 4E (eIF4E) is a rate-limiting component of the eukaryotic translation apparatus. This factor is specifically involved in the mRNA-ribosome binding step of eukaryotic protein synthesis of cap-dependent translation so that the majority of cellular mRNA requires eIF4E to be translated into proteins (Richter JD et al., 2005). In the 48S initiation complex, a role for the eIF4G subunit of eIF4F has also been documented (Grüner S et al., 2016). Specifically, eIF4G is a scaffold protein that binds eIF4E and serves as a central ribosome adaptor module which attracts 40S ribosomal subunits to the 5′ end of mRNAs via direct association with eIF3. We therefore measured the hippocampal levels of the phosphorylated and non-phosphorylated forms of eIF4E, and the levels of eIF4G in hippocampal extracts from 3-month old Tg2576 mice. Results (Fig. 2D-E and Suppl Fig 2) showed that none of these pre-initiation component of translation was altered in this genotype x age condition (p-eIF4E/eIF4E: t_(9)_=0,47 *p* = 0,65; eIF4G: t_(4)_= 0,94 *p* = 0,40).

### Salubrinal reduces hippocampal APP and Aβ 1-42 levels

Salubrinal is a selective inhibitor of cellular complexes that dephosphorylate eIF2α and therefore decreases overall translation. Having shown that APP and Aβ levels are increased at ages where p-eIF2α phosphorylation (Borreca et al., 2016) is reduced, we have examined the possibility that i.c.v. or i.p. injections of the inhibitor of eIF2α dephosphorylation salubrinal could decrease abnormally elevated APP and Aβ levels. Western blot were carried out in hippocampal extracts from 3-month old Tg2576 mice treated with the compound or the vehicle (DMSO). Data are shown in Fig. 3. Statistical comparison of densitometric quantification of APP (Fig. 3A and Suppl Fig 3A) and Aβ (Fig. 3B and Suppl Fig 3A) revealed that these levels were significantly lower in salubrinal-injected mice compared with DMSO-injected mice irrespective of the injection route [APP, i.c.v. t(_10_) = 2,71; *p* = 0.02; i.p. t(_8_)=2,54, *p* = 0.03; Aβ, icv: t_(11)_= 2,20; *p* = 0.049, i.p.: t_(4)_ = 3,33; *p* = 0.03). Of note for potential translational applications, i.p. injections were as much efficient as i.c.v. injections in decreasing hippocampal APP and Aβ levels. This observation prompted us to use the less invasive injection route for subsequent evaluation of the salubrinal rescuing effects on molecular, neural and cognitive alterations found in early symptomatic mutant mice.

**Figure 3:**
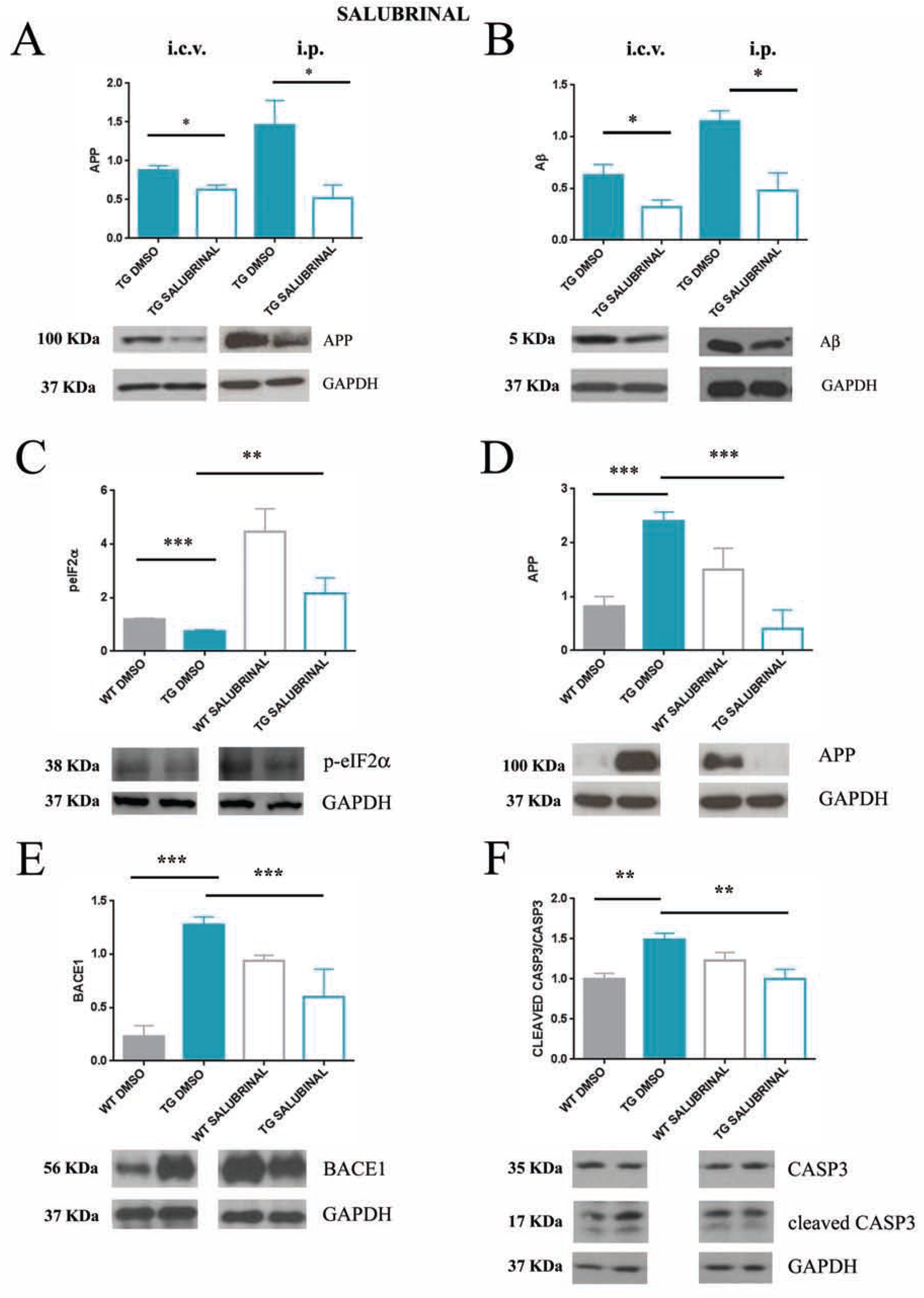
Salubrinal rescues pathogenic AD markers in the hippocampus of 3-month old Tg2576 mice. Histograms showing that i.c.v. (1 μl of a 75 μM solution diluted in DMSO) or i.p. (1 mg/kg diluted in DMSO repeated for 7 consecutive days) injections of salubrinal were effective in decreasing **(A)** APP and **(B)** Aβ levels in the hippocampus of Tg2576 mice. Histograms showing that i.p. injections of salubrinal were effective in restoring proper levels of **(C)** eIF2α, **(D)** APP, **(E)** beta-secretase 1 (BACE-1) in hippocampal extracts, and **(F)** of caspase-3 in hippocampal synaptosomes of 3-month old Tg2576 mice. Bars represent mean ± SEM. (Tg2576 DMSO: solid blue bars; Tg2576 salubrinal: empty blue bars; wild-type DMSO: solid grey bars; wild-type salubrinal: empty grey bars) * *p* < 0,05; ** *p* < 0,01; *** *p* < 0,005).

### Salubrinal restores proper levels of p-eIF2α, APP, BACE1 in hippocampal extracts from early symptomatic Tg2576 mice

The verification that salubrinal rescues early upregulation of translation requires to show that the compound increases p-eIF2α levels of Tg2576 mice up to the levels of WT mice. Results are shown in Fig. 3C and Suppl Fig 3B. Statistical comparisons showed a main effect of genotype (F_(1,10)_=9,93; *p* = 0,01), of treatment (F_(1,10)_=15,77; *p* = 0,003) and of the genotype x treatment interaction (F_(1,10)_=5,21; *p* = 0,04). Post hoc comparisons first confirmed our previous observation in non-treated mice (see Fig. 2) that p-eIF2α levels in the DMSO condition are significantly lower in Tg2576 mice than in WT mice (*p* = 0,02). Consistent with its function of translation initiation inhibitor, salubrinal increased p-eIF2α in both genotypes (salubrinal vs DMSO WT mice, *P* = 0.001; salubrinal vs DMSO Tg2576 mice, *p* = 0.02). Remarkably, eIF2α levels were similar in salubrinal Tg2576 and DMSO WT mice (*p* > 1). For APP, we found a significant genotype x treatment interaction (F_(1,5)=_27.73; *p* = 0,0032). In the DMSO condition, APP levels were higher in Tg2576 mice than in WT mice (*p* = 0,049). Salubrinal decreased APP in the mutant mice (DMSO vs salubrinal Tg2576 mice, *p* = 0,0019) to the level of DMSO-injected mice (salubrinal Tg2576 mice vs DMSO WT mice, *p* = 0,27). BACE1 is the first cleaving enzyme in the APP amyloid pathway. Extensive evidence show that BACE-1 inhibitors decrease Aβ load and reduce neuro-inflammation in fully symptomatic transgenic AD mice (Neumann U et al., 2015). Differently, mixed effects were reported in AD patients especially when these inhibitors were administered peripherally (Georgievska B et al., 2015). Here we show that following peripheral salubrinal treatment, hippocampal BACE-1 levels were reduced in Tg2576 mice and increased in WT mice respectively (Fig. 3E and Suppl Fig 3C). A two-way ANOVA revealed a significant effect of genotype (F_(1,7)_ = 6,15; *p* = 0.042) and of the genotype x treatment interaction (F_(1,7)_ = 79,93, *p* =0.00004). Post hoc comparisons showed that BACE1 levels were higher in DMSO Tg2576 than in DMSO WT mice (*p* =0.0003), and that the treatment was efficient in decreasing the BACE-1 levels of Tg2576 mice to those of DMSO WT mice (*p* =0,42).

### Salubrinal rescues AD-selective caspase-3 alteration in hippocampal synaptosomes from early symptomatic Tg2576 mice

Caspase-3 (Casp-3) activity is significantly increased at hippocampal synapses of Tg2576 mice (D’Amelio M et al., 2011) where it causes permanent activation of calcineurin which leads to dephosphorylation of the AMPA receptor GluR1 subunit and its removal from postsynaptic sites (D’Amelio M et al., 2011). In association with these indexes of synaptic failure, mice show a reduction in CA1 spine density, a decrease in basal glutamatergic synaptic transmission associated with an increase in long-term depression (LTD) at CA1-CA3 synapses, and impairments in hippocampal dependent memory. Because protein synthesis is required for Casp-3 activation (Coxon FP et al., 1998), we examined the possibility that salubrinal-mediated decrease of eIF2α phosphorylation, which reduces protein translation, could normalize Casp-3 levels in the mutant mice. Results are shown in Fig. 3F and Suppl Fig 3D. In line with this hypothesis, statistical comparison of the cCasp-3/Casp-3 ratio measured in the four mice groups revealed a significant genotype x treatment interaction (F_(1,8)_ = 1,13 *p* = 0.004). Pair comparisons then showed that these levels were reduced in salubrinal Tg2576 mice compared to DMSO Tg2576 mice (*P* = 0.006) and were not different from those detected in DMSO WT mice (p =0.99). To our knowledge, these data provide the first *in vivo* demonstration that, as *in vitro* (Gong N et al., 2015), salubrinal is efficient in decreasing apoptosis markers at synapses.

### Effect of salubrinal in WT mice

Consistent with the report (Mouton-Liger F et al., 2012) that increasing eIF2α phosphorylation in healthy mice increase BACE-1 levels and induces Aβ amyloidogenesis, higher levels of BACE-1 (*p* = 0.0006) (Fig. 3E) and p-eIF2α (*p* = 0.001) (Fig. 3C) were detected in the WT mice treated with salubrinal compared to the WT mice treated with DMSO.

### Salubrinal rescues synaptic plasticity alterations

The synaptic plasticity deficit shown in 3-month old Tg2576 mice consists in increased magnitude of CA1-to-CA3 DHPG-induced long-term depression (LTD) (D’Amelio M et al., 2011). We applied the same LTD induction protocol in acute hippocampal slices obtained from Tg2576 and WT mice treated with salubrinal or DMSO. The data are shown in Fig. 4 A-C. A TWO-WAY ANOVA carried out on relative changes of fEPSP slopes before and after low frequency stimulation in DMSO- and salubrinal-injected Tg2576 (Fig. 4A) and WT (Fig. 4B) mice revealed a significant genotype x treatment interaction (F_(1,35)_ = 16.87; *P* < 0.01). Pair comparisons of LTD percentage data (Fig. 4C) showed that DMSO-injected Tg2576 mice had a significant increase in the magnitude of CA3-to-CA1 LTD compared to DMSO-injected WT mice (*p* < 0,001) which was decreased by salubrinal treatment (salubrinal Tg2576 vs DMSO Tg2576 mice, *p* < 0,001) to the value observed in DMSO-treated WT mice (salubrinal Tg2576 mice vs DMSO WT mice, *p* > 1).

**Figure 4:**
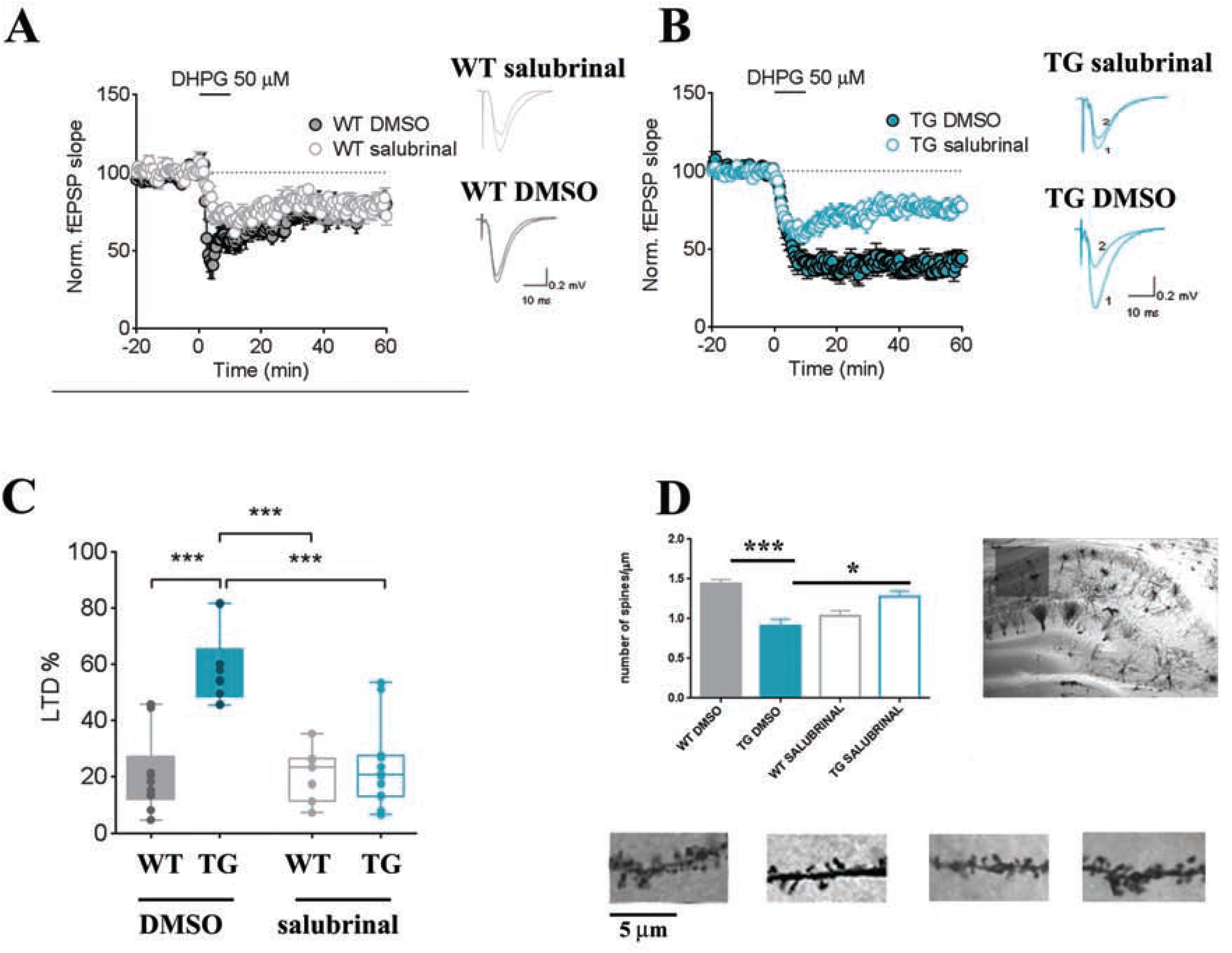
Salubrinal rescues hippocampal CA1 dendritic spine loss and increased LTD at CA3-CA31 hippocampal synapses. Superimposed fEPSPs traces recorded immediately before DHPG (50 μM, 10 min) and at 55-60 min of DHPG washout in 3-month-old saline and salubrinal-treated WT **(A**) and Tg2576 (TG) mice **(B).** Running plots show normalized fEPSP mean slope (± SEM. displayed every 1.5 min) recorded from the dendritic region of CA1 neurons in hippocampal slices from WT and TG mice exposed to 50 μM DHPG for 10 min. **(C)** The box-and-whisker plot indicates the degree of DHPG-LTD, measured as fEPSP slope decrease from baseline, 55–60 min from DHPG washout (WT: n = 10 slices from 4 saline, 7 slices from 4 salubrinal-treated mice; Tg2576: n = 10 slices from 4 saline, 12 slices from 4 salubrinal-treated mice). *** *p* < 0,005 **(D)** Left: Histograms showing dendritic spines (number of spines/dendrite segment length) counted on 5 segments per neuron in 10 pyramidal neurons laying in the CA1 subfield of the hippocampus in wild-type mice (grey bars) and Tg2576 mice (blue bars) injected with DMSO (solid bars) or salubrinal (empty bars). Bars represent mean ± SEM. Right: representative images of Golgi-stained sections of the dorsal hippocampus (scale bar: 10 μm) and of apical dendrite segments of CA1 hippocampal pyramidal neurons (scale bar: 10 μm) in 3-month-old wild-type and Tg2576 mice injected with DMSO or salubrinal. *** *p* < 0,001

Dendritic spines are among the first synaptic elements which are disrupted during cognitive decline (Scheff SW et al., 2007). Consistent with these findings, we found that spine density was significantly decreased in CA1 pyramidal neurons already in 3-month old Tg2576 mice (D’Amelio M et al., 2011). Whether AD-associated early defects in protein synthesis are involved in this effect has not been yet investigated although it could be implicitly assumed that spine loss might be the consequence of a global decrease in protein synthesis. Conversely, we found that salubrinal, which blocks protein synthesis was effective in rescuing spine defects thereby suggesting that the treatment was overall preventing the accumulation of toxic products (Fig.. 4D). Statistical analysis of spine density scores revealed a significant genotype x treatment interaction (F_(1,10)_=27.37; *p* = 0.0038). Pair comparisons then showed that spines were increased Tg2576 mice following salubrinal treatment (*p* =0.02) and that the spines scores were found in salubrinal Tg2576 mice and in WT mice irrespective of the treatment (*P* > 1 for each comparison). It is therefore apparent that there is an early phase in which enhanced APP levels are harmful for hippocampal spines and that salubrinal-mediated decrease of APP levels exerts synapse-protective effects.

### Salubrinal rescues Novel Object Recognition performance and neuronal activity

Early synaptic deficits shown by Tg2576 mice correlate with impairments in hippocampal-dependent tasks. Accordingly, we probed whether rescuing the plasticity and morphology of hippocampal synapses also rescued novel object recognition performance. The data are shown in Fig. 5. During the training phase (Fig. 5B), no difference was observed in the rate of exploration of the two identical objects in any group (DMSO WT: object sx vs object dx, t_(10)_=0,10; p = 0,9; DMSO Tg2576: object sx vs object dx, t_(8)_=1,66 p = 0.13; salubrinal WT: object sx vs object dx, t_(14)_=0,84; p = 0,42; salubrinal Tg2576: object sx vs object dx t_(10)_=1,87; p = 0,09) while, during the test phase (Fig. 5C) only DMSO Tg2576 mice failed to explore more the novel (NO) than the familiar (FO) object (DMSO Tg2576: NO vs FO, t_(8)_=0,29 p = 0,77; DMSO WT: NO vs FO t_(12)_=3,86; p = 0,0022; salubrinal WT: NO vs FO t_(16)_=2,18; p = 0,04; salubrinal Tg2576: NO vs FO, t_(18)_=3,65; p = 0,0018).

Based on data showing that Tg2576 mice show defective activation of hippocampal neurons during formation or retrieval of hippocampal-dependent memory (Broadbent NJ et al., 2004; Lelos MJ et al., 2014), and that healthy mice exposed to the NOR test show hippocampal neuronal activation (Tanimizu T et al., 2017), we visualized by immunofluorescence techniques the protein product of the *c-fos* proto-oncogene in the CA1 region 1 h after completion of the NOR test. The data are shown in Fig. 5D. A two-way ANOVA carried out on the number of immunoreactivity spots detected in the CA1 region of the four experimental groups revealed a significant genotype x treatment interaction (F_(1,22)_= 8,04; *p* = 0.009). Pair comparisons then showed a lower number of *c-fos* immunoreactive spots in DMSO Tg2576 mice compared with DMSO WT mice (*p* = 0.0002) that was rescued that salubrinal treatment (salubrinal Tg2576 vs DMSO WT mice, *P* = 0,76). No effect of treatment was detected on *c-fos* expression in the wild-type mice (salubrinal WT vs DMSO WT *p* = 0,83).

## DISCUSSION

Accumulation of Aβ peptides resulting from amyloidogenic APP cleavage is strongly involved in AD and other intellectually disabling diseases including Down syndrome or Fragile-X syndrome. Hence, strategies for treating these diseases have massively focused on designing drugs inhibiting the production or facilitating the clearance of Aβ fragments or, alternatively, producing genetic (Hamilton A et al., 2014) or pharmacological (Hamilton A et al., 2016) inactivation of the metabotropic glutamate receptor 5 (mGluR5) which acts as a receptor for Aβ and mediates synaptic dysfunction. Surprisingly, despite evidence showing that full length APP is over-expressed in the AD brain (Johnston JA et al., 1994), and that downregulation of APP levels reduces Aβ deposition in APP751SW cells (Asuni AA et al., 2014) and in the brain of symptomatic AD mice (Lahiri DK et al., 2007; Asuni AA et al., 2014; Teich AF et al., 2017), the idea that regulation of APP levels *per se* might be beneficial for contrasting AD has been poorly considered.

APP overexpression is the result of upregulation of APP mRNA translation. Regarding Tg2576 hAPP mutants, without excluding the presence of specific translational influences deriving from the hamster prion promoter, there is evidence that once the transgene is inserted in the mouse genome, the regulators of hAPP translation in the symptomatic phase undergo the same alterations as those observed in human patients. For example, consistent with data showing that the two RNA binding proteins, Fragile-X Mental Retardation Protein (FMRP) and heteronuclear Ribonucleoprotein C (hnRNP C), exert an opposite control on APP translation (Lee EK et al., 2010; Westmark CJ et al., 2012), we reported that the enhancement of hAPP levels in hippocampal extracts from both symptomatic Tg2565 mice and sporadic AD patients is accompanied by a downregulation of FMRP and an upregulation of hn RNP C (Borreca et al., 2016). We also noticed in the mutant mice that the stronger increase in hAPP and the maximal dysregulation of FMRP and hnRNP C were observed when mice were 1- and 3- month old, i.e., before or immediately after Aβ oligomers, synaptic failure and cognitive deterioration could be detected. These data prompted us to examine the link between longitudinal variations in hAPP expression and dysregulation of specific *vs* overall translation.

A polyribosome profiling analyses of hAPP in total hippocampal extracts from Tg2576 mice and WT mice first showed that the presence of the protein in polysomal fractions was exclusively detected when mice were 1- and 3-month old. hAPP mRNA was also more expressed in polysomal fractions at any age, but a peak of expression was detected at the 3-month age point concurrently with the release of Aβ oligomers. For overall translation, we measured the levels of several eukaryotic initial translation factors and found a significant decrease of the p-eIF2α/eIF2α ratio in 1- month and 3-month mutant mice compared to WT mice, which indicates that translation was upregulated at stages where hAPP undergoes maximal expression. Differently, confirming that the translational machinery is downregulated in association with the full symptomatology, no hAPP mRNA signal was detected in symptomatic mice and sporadic AD patients but the p-eIF2α/eIF2α ratio was found to be increased in 6- and 9-month old Tg2576 mice, consistently with previous data from symptomatic mice (Kim HS et al., 2007) and AD patients (Ma et al., 2013). A plausible explanation for this phenomenon might be that initial low levels of p-eIF2α increase protein synthesis, lead to APP accumulation, disrupt physiological APP cleavage, promote Aβ accumulation and determine, in turn, microglia activation and oxidative stress (Hetz C et al., 2017). Thus, the late increase of eIF2α phosphorylation might act as a compensatory/neuroprotective mechanism aimed at restoring physiological APP metabolism via a downregulation of overall translation. Interestingly, we found no evidence of alteration of other main players of translation initiation including the phosphorylated and non-phosphorylated states of the eukaryotic translation initiation factor 4E (p-eIF4E and eIF4E), and the non-phosphorylated state of the eukaryotic translation initiation factor 4G (eIF4G), suggesting a highly selective role for p-eIF2α in mediating AD-specific dysregulation of translation. It is therefore apparent that the phosphorylation status of eIF2α varies according to the progression of AD pathogenesis with an initial unbalance in favor of decreased phosphorylation, compatible with enhanced translation, followed by a late unbalance in favor of increased phosphorylation, compatible with decreased translation (Table 1).

**Table 1.** Variations in p-eIF2α levels reported in TG mice and AD patients. If detected in tissues from patients with mild cognitive impairment, the opposite dysregulation of p-eIF2α and APP levels might provide an ultra-early AD marker.

|  | early symptomatic | symptomatic |
| --- | --- | --- |
| mice | APP ↑<br>p-eIF2α ↓<br>(present study) | APP ↑<br>p-eIF2α ↑<br>[Kim et al., 2007] |
| human patients | APP ↑<br>p-eIF2α ? | APP ↑<br>p-eIF2α ↑<br>(Chang et al., 2002<br>Ma et al., 2013) |

Phosphorylation of the α-subunit of eukaryotic initiation factor 2 (eIF2α) acts as a negative regulator of general translation (Donnelly N et al., 2013; 36 Ohno M., 2014). In AD, there is evidence that accumulation of Aβ increases eIF2α phosphorylation which then blocks mRNA translation and *de novo* protein synthesis. This has been shown in cultured cells expressing swe-hAPP and in brain tissues from sporadic AD patients (Kim HS et al., 2007; Chang RC et al., 2002; Segev Y et al., 2013) or from symptomatic AD mice expressing the hAPP mutation alone (Kim HS et al., 2007) or in association with other mutant proteins (Devi L et al.,2010; Devi L et al., 2010a; Devi L et al., 2013; Page G et al.,2006). Accordingly, enhancement of p-eIF2α has been considered as the mechanism which mediates cognitive deterioration by downregulating expression of synaptic plasticity proteins. More recently, however, elevated levels of eIF2α phosphorylation have been shown to associate with increased translation of a subset of mRNAs among which the β-secretase enzyme BACE-1, one main determinant of amyloidogenic APP cleavage (Devi L et al., 2010; O’Connor T et al., 2008). This observation, together with the finding that activation of peIF2α elicits the unfolded protein response in the hippocampus and the temporal cortex of AD patients (Hoozemans JJ et al., 2005) reveal that downregulation of translation activates prominent compensatory mechanisms aimed at restoring normal cell function.

Having identified an early symptomatic age point (3 months) wherein eIF2α is decreased, hAPP and hAPP mRNA are increased, and Aβ oligomers become detectable, we then examined whether inhibiting eIF2α dephosphorylation, i.e., decreasing overall translation, could prevent the elevation of hAPP and Aβ levels. We tested this hypothesis by injecting the selective inhibitor of eIF2α dephosphorylation salubrinal both i.c.v. and i.p. in 3-month old mutant and found that, whatever the injection route, salubrinal significantly lowered hippocampal APP and Aβ levels. Our findings are in line with *in vitro* data showing that short-term treatment with salubrinal attenuates Aβ-induced neuronal death in primary cortical neuronal cells (Huang X et al., 2011), and with several *in vitro* and *in vivo* data showing that compounds like MMP13 which produces translation regulation of BACE-1 (Zhu et al., 2019), or posiphen, which inhibits APP translation, (Teich AF et al., 2018) decreases the production of toxic Aβ, and rescues cognitive deficits

We therefore verified whether salubrinal administered via the less invasive intraperitoneal injection route could restore physiological levels of those and other major pathogenic AD markers, and prevent the manifestation of neural and cognitive symptoms. We first observed that hippocampal levels of APP, Aβ, BACE-1 measured in salubrinal-injected Tg2576 mice no longer differed from those measured in vehicle-injected WT mice. In the mutant mice, peripheral salubrinal administration also abolished the enhancement of caspase-3 and the prevalence of LTD at hippocampal synapses, prevented dendritic spine loss in pyramidal hippocampal neurons, rescued performance impairments in the hippocampus-dependent NOR task, and restored NOR-induced hippocampal *c-fos* activation.

Altogether, our findings show an early upregulation, followed by a downregulation, of translational efficiency which, by favoring haPP overexpression could precipitate the transition between the prodromal and the symptomatic phase. Remarkably, repressing overall translation by salubrinal at the onset of AD-like symptoms does not only prevent the release of toxic products (Aβ) or apoptotic factors (caspase-3), but reinstates APP physiological processing via concurrent regularization of APP and BACE-1 levels. Considering that abnormally elevated amounts of APP to be cleaved are expected to saturate α-secretase function, dysregulate physiological (non-amyloidogenic) processing, and favor the emergence of pathological (amyloidogenic) processing, it is therefore tempting to hypothesize that prodromal upregulation of translational efficiency plays a critical role in triggering AD pathology. Thus, while our findings confirm that overall translation is decreased in symptomatic AD, they reveal that it is transiently enhanced before the manifestation of full AD symptoms suggesting, in turn, that concurrent enhancement of APP and eIF2a levels at the first signs of cognitive deterioration should reliably predict future AD.

## ABBREVIATIONS

eIF2α: Eukaryotic Initiation Factor 2
BACE1: Beta-secretase 1
hAPP: human amyloid precursor protein
FMRP: Fragile X Mental Retardation Protein
hnRNPC: Heterogeneous Nuclear Ribonucleoprotein C (C1/C2)
DMSO: Dimethyl sulfoxide
CASP-3: caspase-3
GDP: Guanosin Diphosphate
GTP: Guanosine Tiphosphate
eIF4G: Eukaryotic translation initiation factor 4 G
eIF4E: Eukaryotic translation initiation factor 4E
DHPG: 

## ACKNOWLEDGEMENTS

This work was supported by a grant (AGESPAN) from CNR-National Research Council of Italy to M A-T. A.B. is currently supported by Fondazione Umberto Veronesi. MDA was supported by the Italian Ministry of Health (Progetto Giovani Ricercatori Project Code GR-2011-02351457) and by a grant from the Alzheimer’s Association, United States (Project Code AARG-18-566270).

We thank Marcello Ceci for technical support with polysome gradient machinery and Stefano Biffo for scientific support and constructive reading of the manuscript

## Authors contribution

A.B. conceived and designed the research plan, supervised the molecular, anatomical, and behavioral experiments, and wrote the manuscript. F.V., L.E., M.D.L. carried out polysome profiling, western blot, behavioral, anatomical, and c-fos experiments. A.R. performed eIF4G and eIF4E analyses. A.C. run the electrophysiological experiments under the N.B.M supervision. V.C. and GA performed p-eIF2α measurements in 3-month old mice. A.N. performed caspase-3 analysis in hippocampal synaptosomes under the M.D.A. supervision. M.D.A. critically read the manuscript. M.A.T. supervised the research plan and wrote the manuscript.

## Supplementary

**Supplementary Figure 1**: Histograms showing the distribution of hAPP mRNA on P and NP fractions after further processing of total RNA from each fraction by RT–qPCR (normalization with the synthetic RNA Kanamicin) in post-mortem hippocampal samples from two sporadic AD patients.

**Supplementary Figure 2**: Total blot of Figure 2 measurements of p-eIF2α (A-B), eIF4E (C), eIF4G (D).. GAPDH was used for normalization.

**Supplementary Figure 3**: Total blot of Figure 3 measurements of APP, Aβ, BACE-1 and caspase- 3 levels. GAPDH was used for normalization.

